# Trying harder: how cognitive effort sculpts neural representations during working memory

**DOI:** 10.1101/2023.12.07.570686

**Authors:** Sarah L. Master, Shanshan Li, Clayton E. Curtis

**Affiliations:** Department of Psychology, New York University; Program in Psychology, New York University Abu Dhabi; Center for Neural Science, New York University

**Keywords:** working memory, cognitive effort, fMRI, modeling, human, retinotopy, decoding

## Abstract

The neural mechanisms by which motivational factors influence cognition remain unknown. Using fMRI, we tested how cognitive effort impacts working memory (WM). Participants were precued whether WM difficulty would be hard or easy. Hard trials demanded more effort as a later decision required finer mnemonic precision. Behaviorally, pupil size was larger and response times were slower on hard trials suggesting our manipulation of effort succeeded. Neurally, we observed robust persistent activity in prefrontal cortex, especially during hard trials. We found strong decoding of location in visual cortex, where accuracy was higher on hard trials. Connecting these across-region effects, we found that the amplitude of delay period activity in frontal cortex predicted decoded accuracy in visual cortex on a trial-wise basis. We conclude that the gain of persistent activity in frontal cortex may be the source of effort-related feedback signals that improve the quality of WM representations stored in visual cortex.

## Introduction

Undoubtedly, motivation and cognition interact. Increases in motivation, both extrinsic and intrinsic, often improve higher-order cognitive performance^1–3^. The exertion of cognitive effort is one possible mechanism by which motivation could improve cognitive performance^4–6^. Here, we define cognitive effort as the exertion of a general cognitive resource in the service of improving performance on a task. This resource can be exerted in response to extrinsic (e.g., rewards) or intrinsic (e.g., drive to succeed) motivational factors^7^. The interaction of motivational and cognitive variables makes it difficult to isolate the influence of one or the other on cognitive performance, particularly in psychiatric conditions, where motivational and cognitive dysfunction are often comorbid^8–10^. Isolating the impact of cognitive effort on higher-order cognition, with other motivational variables aside, is the first step in the innovation of targeted interventions to ameliorate the cognitive deficits which accompany psychiatric disorders.

Working memory (WM) supports the online maintenance and manipulation of information^11^. As such, WM is a necessary building block for most high-level cognitive abilities including navigation, learning, and decision-making^12–14^. However, the capacity of WM storage has severe limitations^15–18^, so the optimal use of WM requires strategic allocation of resources based on task demands and potential rewards. Indeed, humans can flexibly allocate their WM resources and improve WM accuracy according to the priority and incentives of memoranda^1,19–21^. What remains unknown is how the exertion of cognitive effort might impact WM performance, and the neural mechanisms by which this occurs. Effort may facilitate WM performance through an increased recruitment of neural resources and/or the gain of neural activity. Indeed, previous work has demonstrated increased fMRI activity in prefrontal cortex as WM load increases^22,23^, and we suggest this increase could have been mediated in part by increased effort.

Here, we hypothesized that effort facilitates more accurate WM by preventing drift of mnemonic representations during the WM delay (i.e., reducing memory error), and/or by reducing noise in representations (i.e., reducing memory uncertainty). To test this hypothesis, we measured fMRI activity from 12 humans performing a visuospatial WM task that varied the effort required for success. As predicted and consistent with past work, we found that cognitive effort was associated with increased pupil size^24,25^, response times^26–29^, and amplitudes of delay period activity in frontal and parietal cortex^22,30^. Critically, using Bayesian decoding^31,32^ we found that WM representations decoded from early visual cortex were more accurate on more effortful trials. To potentially link these effects, we hypothesized that effort, encoded in the amplitude of persistent activity in frontal cortex, impacts the quality of WM representations encoded in visual cortex. Indeed, persistent activity in frontal cortex predicted decoding in visual cortex on a trial-by-trial basis. These results indicate that the exertion of cognitive effort improves WM performance. They suggest that the exertion of effort results in diffuse increases in neural gain, which serve to stabilize WM representations and prevent memory drift. Moreover, the fidelity of WM increases associated with cognitive effort may be mediated by feedback signals from frontal cortex that sculpt population activity in visual cortex.

## Results

We measured fMRI activity while participants performed a visuospatial WM task that differed in the degree of cognitive effort required. We operationalized cognitive effort by manipulating the difficulty of a judgment to be made at the conclusion of each trial (Figure 1A). At the start of each trial, we presented a cue which indicated the difficulty of the decision after the memory delay, in which participants judged whether a probe was clockwise/counterclockwise of a memorized target. One psychophysical staircase (90% target accuracy) ensured that easy trials could be successfully performed with less memory precision and thus less effort. Another staircase (70% target accuracy) ensured that hard trials required more precision and thus more effort. These staircases resulted in an average distance of 23 degrees of visual angle between the target and the test probe on easy trials. This distance was only 5 degrees on hard trials.

**Figure 1.**
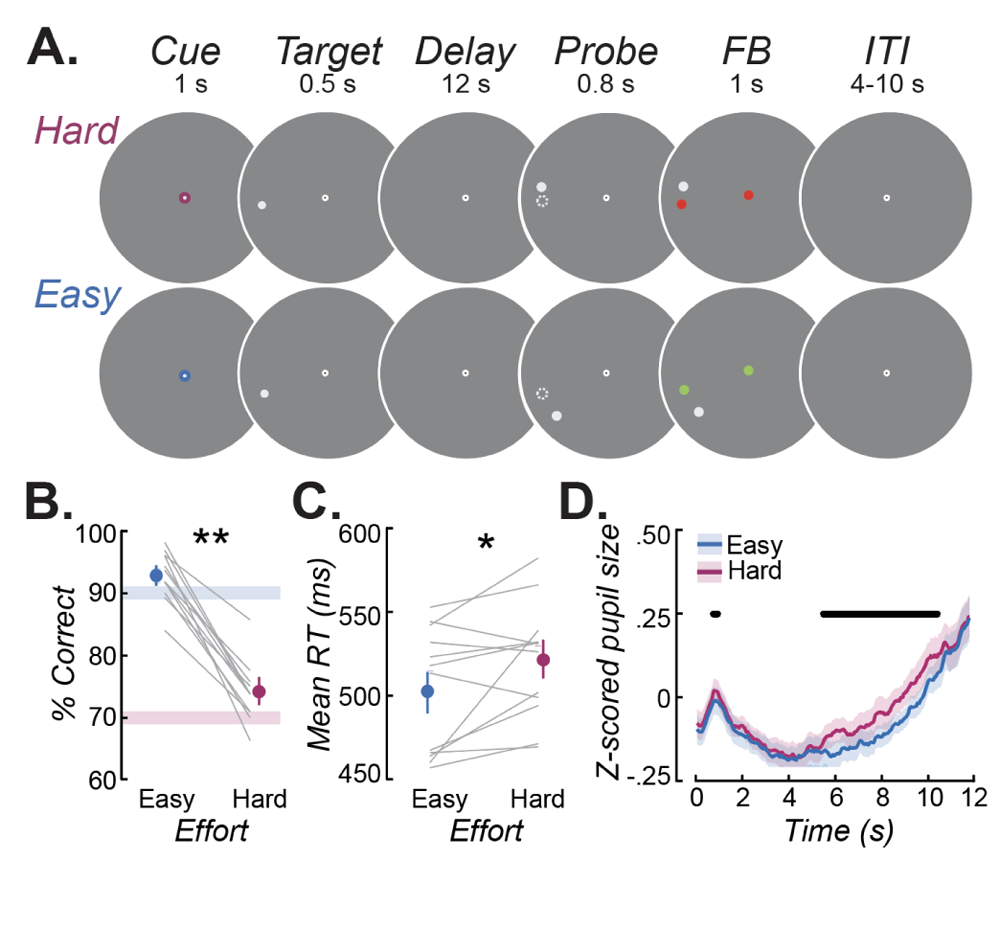
Task design and behavioral results. **A.** There were two interleaved types of spatial WM trials that differed in the degree of cognitive effort required for successful performance. At the beginning of each trial the color of the fixation point indicated whether the discrimination after the memory delay required coarse (easy) or fine (hard) mnemonic precision. Participants maintained the location of a briefly presented target over a long memory delay. Then, they indicated with a button press whether a probe stimulus (dot) was clockwise or counter clockwise to the memorized target location. In the example above, the correct answer is counter clockwise for the hard and clockwise for the easy trial. After the choice, the fixation and the re-presented target dots turned green (correct) or red (incorrect) as feedback (FB). **B.** Controlled by two staircase procedures, participants were more accurate on easy compared to hard trials. The blue and pink lines mark the target accuracies for each staircase. **C.** Response times (RT) were longer for hard compared to easy trials. Gray lines are individual participants and dots are group means. Error bars are standard errors of the mean. **D.** Pupil sizes were larger during the delay period for hard compared to easy trials. Lines depict the mean, while error bars are standard errors of the mean. Black lines indicate significant differences in pupil dilation between hard and easy trials Bonferroni-corrected p<0.00016. These behavioral data confirmed that our manipulation of effort was successful.

We then evaluated whether subjects exerted more effort during the WM delay on hard trials, in anticipation of the difficult upcoming judgment. We measured two well-known correlates of cognitive effort, pupil size^24,25,33^ and reaction time^26–29^. As expected from the staircase procedure, participants were more accurate on easy compared to hard trials (p<0.001; Figure 1B). They were also slower to respond on hard than easy trials (p<0.05; Figure 1C). Moreover, pupil size during the delay period was larger during hard compared to easy trials. This effect was significant at the group level from seconds 7 to 12 of the delay period (all p’s<0.00016; Figure 1D), and within 9 of the 12 participants when considering the entire delay period in (confidence intervals on regression calculated with *α*=0.05; see Methods). Overall, the behavioral data confirmed the effectiveness of our manipulation of effort.

We then asked whether cognitive effort impacted the gain of persistent activity in multiple retinotopically defined^34^ visual field maps (Figure 2A; see *Retinotopic Mapping Task* and *pRF Model Fitting and ROI drawing* in Methods). We plotted BOLD activity averaged over hard and easy trials separately (Figure 2B). Several patterns emerged from these data. First, following the target-evoked phasic response seen across all maps, activity persisted throughout the delay period. The amplitude of persistent activity increased up the visual hierarchy. We found the strongest persistent activity in frontal cortex, while in visual cortex the visually evoked response returned to pre-trial baseline levels. Additionally, persistent activity was higher when the memory target was in the voxels’ receptive fields (“RF in”) compared to when it was in the opposite part of the visual field (“RF out”; note the differences between solid lines and dashed lines). Finally and central to our main hypothesis, persistent activity was higher during hard compared to easy trials (note the differences between magenta and blue lines). We quantified these effects using a 3-way, repeated measures ANOVA on mean late delay period (seconds 6-12) BOLD activity (gray epoch in Figure 2B). The main effects of trial difficulty (hard/easy; F=22, p<0.001), voxel RF (in/out; F=24, p<0.001), and ROI (F=27, p<0.0001) were all significant (permutation testing, p<0.05). The interaction between trial difficulty and ROI (F=8, p<0.0001) was significant, but other interactions between ROI and RF (F=1.7, p>0.1) and RF and trial difficulty (F=1.3, p>0.1) were not. We followed up on the interaction between ROI and difficulty with post-hoc t-tests. Hard trials elicited significantly greater BOLD activity overall (p<0.01). Within ROIs, this effect was strongest within the precentral sulcus (V1-V2-V3 p>0.1, V3AB p>0.1, IPS0-IPS1 p>0.1, IPS2-IPS3 p=0.08, iPCS p<0.05, sPCS p=0.06). Although these results indicate that the exertion of cognitive effort increased BOLD responses, leveraging the pRF properties further show that effort had an equal effect on voxels across the visual field map regardless of whether they contained the memorized target (there was no interaction between trial type (hard/easy) and RF type (in/out) on BOLD activity). This suggests that the application of cognitive effort evokes diffuse activation gain over the entire topographic map, not just the task-relevant portion of the map.

**Figure 2.**
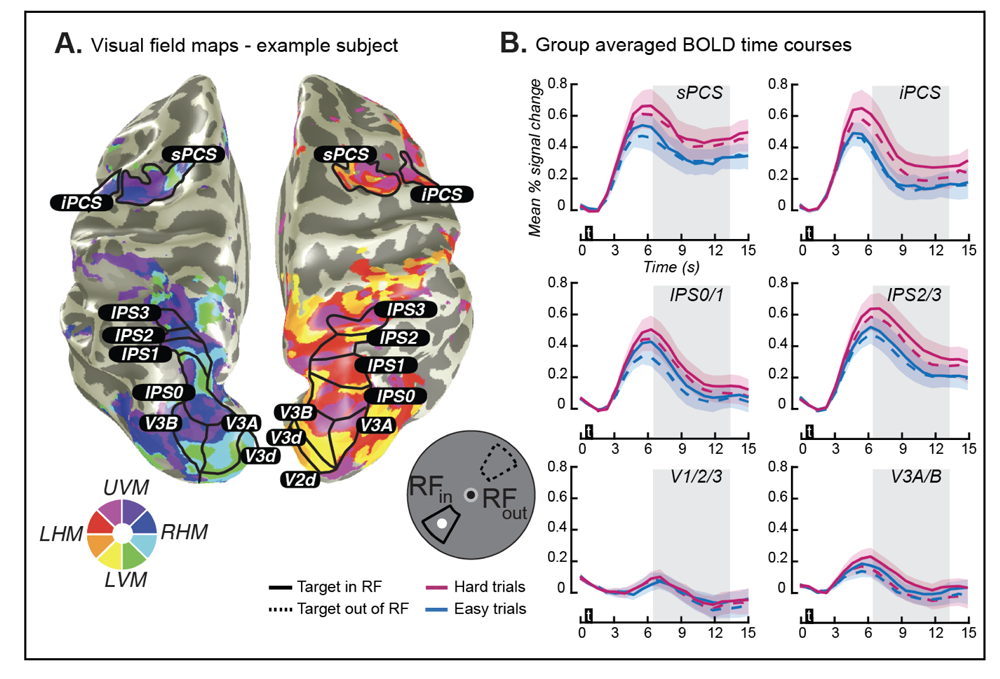
Visual field maps and trial average BOLD time courses. **A.** Polar angle maps derived from population receptive field mapping were used to define visual field map ROIs in each subject. **B.** BOLD time courses averaged across hard and easy trials. Solid lines denote mean BOLD signal when the target was within 15 degrees of the voxel’s receptive field center. Dashed lines denote mean BOLD signal when the target was within 15 degrees opposite the voxel’s receptive field. Error bars are the standard error of the mean computed across participants. Black boxes with “t” inside indicate when the target was present. The gray boxes indicate the epoch used for statistical comparisons of BOLD activity and used as input to the Bayesian decoder.

Next, we aimed to test if cognitive effort had an effect on the quality of WM representations. We used a Bayesian decoder (see *Methods*) to estimate the accuracy and uncertainty of WM representations^35^ as either of these metrics of the memory quality could be affected by effort. We predicted that effort improves the accuracy of WM representations by reducing the error between decoded and true memorized locations, and therefore decoded error would be smaller on hard compared to easy trials (Figure 3A). Similarly, we predicted that effort reduces the uncertainty of WM representations estimated from the widths of decoded probability distributions over memorized locations. We reasoned that hard trials would have less uncertainty than easy trials. For illustrative purposes, we plotted two example trials, one hard and one easy, from V3A/B in one participant (Figure 3B). Note that in this example the hard and easy trials differed only in terms of accuracy (error between decoded and real target location) and not uncertainty (width of each decoded probability distribution). In fact, these two example trials are representative of what we found across trials and participants in visual and posterior parietal cortex. Effort impacted the accuracy (Figure 3C) and not uncertainty (Figure 3D) of memorized representations. Two-way repeated measures ANOVAs on decoded error and decoded uncertainty yielded a main effect of trial difficulty (hard/easy) on decoded error, but not on decoded uncertainty (error F=14, p<0.01; uncertainty F=0.05, p>0.05). Following up on this main effect, we found that decoded error was lower on hard compared to easy trials in ROIs V1-V2-V3 (p<0.01), V3AB (p<0.01), and IPS0-IPS1 (p<0.05). There was no difference in decoded accuracy between hard and easy trials in IPS2-IPS3, iPCS, or sPCS, which was not surprising as we found poor or non-existent decoding in these areas when we ignored trial type (hard/easy) (Supplementary Figure 1). In summary, we found that the exertion of cognitive effort resulted in more accurate decoding from visual and parietal cortex.

**Figure 3.**
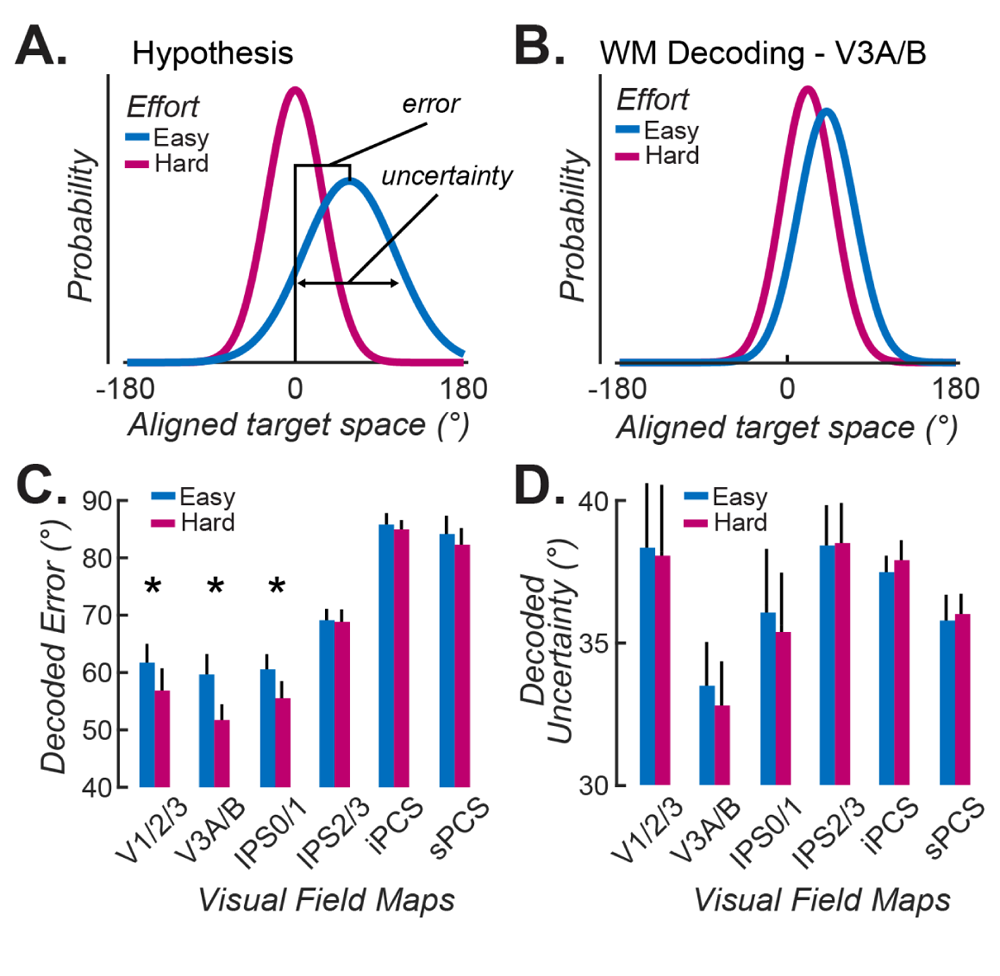
Cognitive effort impacts the accuracy but not uncertainty of decoded WM representations. **A.** Hypothetically, effort could impact the accuracy of WM representations, which would be reflected in decoded error, the difference between the true and decoded location. Effort could also impact the uncertainty of WM, reflected in the width of the decoded WM representation. **B.** Decoded probability distributions for hard and easy trials from area V3A/B in an example subject, but consistent with our group results. Note that effort only impacted the decoded accuracy and not uncertainty of WM, as only the error and not width of the distribution was affected by trial type. **C.** Decoded memory error in early visual cortex (V1/2/3), V3A/B, and IPS0/1, but not more anterior areas, was worse for Easy compared to Hard trials. Bars are means across participants, and error bars are standard errors of the mean. * indicates p<0.05. **D.** Decoded memory uncertainty did not differ between hard and easy trials.

Finally, we hypothesized that the increased gain of persistent activity in frontal and parietal cortex associated with cognitive effort might be the source of feedback control signals that target early visual cortex. We predicted that these feedback signals impact the quality of WM representations. Recall, we could not decode the content of WM from frontal and parietal cortex, despite the fact that these areas showed robust persistent activity during the memory delay. Early visual cortex expressed the opposite pattern to frontoparietal cortex; despite very little evidence of persistent activity, we could decode the locations of memoranda with precision.

To identify potential sources of top-down control over the quality of WM representations stored in early visual cortex, for each participant we computed correlations between decoded WM error from activity patterns in V1-V2-V3 on each trial and the magnitude of delay period activity on each trial for every voxel in the brain. Such a correlation identifies voxels whose increases in gain of persistent activity predicted decreases in decoded error of WM in early visual cortex. Because effort had an impact on decoded error, we focus on this metric of representational quality, rather than uncertainty (note that we found little evidence that persistent activity related to decoded uncertainty; Supplementary Figure 2). Remarkably, we observed large clusters of negative correlations in the superior precentral sulcus and along the intraparietal sulcus that survived conservative cluster based corrections for multiple comparisons (Figure 4A; Methods). In order to estimate if these clusters belong to retinotopic areas in the precentral and intraparietal sulci, we plotted each participants’ sPCS, IPS0/1, and IPS2/3 maps from their native to standard atlas space. We quantified overlap by the number of participants whose maps were co-registered on the cortical surface (Figure 4B). The clusters of correlation localized to the sPCS and IPS2/3 maps most clearly. To summarize, the areas which showed strong persistent activity that was greater when more effort was exerted (i.e. sPCS, IPS0-1, and IPS2-3) were identified by these correlations as possible sources of control over WM representational quality in V1-V2-V3.

**Figure 4:**
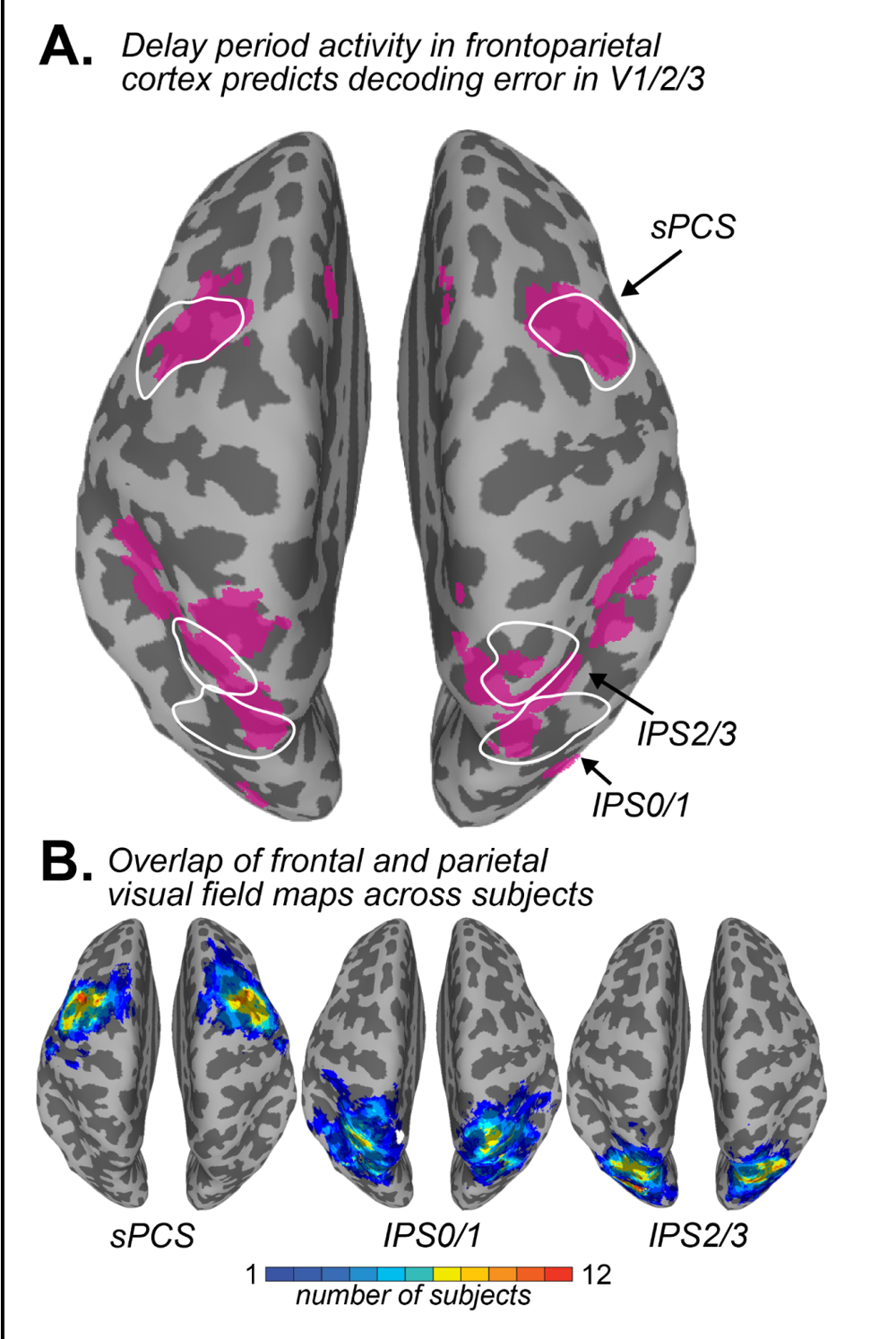
Gain of delay period activity predicts decoded error from early visual cortex. **A.** To identify potential sources of top-down control over representational fidelity in visual cortex, we created a whole-brain correlation map. Trial-by-trial magnitudes of delay period activity across the whole brain were the predictors. The predicted term was trial-by-trial decoded error from V1-V2-V3. The magenta clusters are areas with significant negative correlation at the group-level using cluster-based permutation testing. A negative correlation with decoded error indicates a positive relationship between representational quality in visual cortex and delay period magnitude (magenta). **B.** Single subject ROI overlap in MNI space (sPCS, IPS0/1 and IPS2/3). The color of the heatmap indicates the number of participants whose ROIs overlapped on the cortical surface. The hotter colors (orange, red) indicate that more participants had overlapping ROIs at that location, and the cooler colors (green, blue) very little overlap between participants. Areas with overlap of at least 50% of subjects are drawn in white contours on A.

## Discussion

Despite its importance, understanding how effort impacts cognition remains challenging because the construct of effort is difficult to precisely define and operationalize^4,36,37^. Here, we relied on the simple intuition that some tasks require more effort than others and success depends on how hard we try. We tested the hypothesis that the exertion of cognitive effort improves the quality of WM representations encoded in the brain. We predicted that participants would exert more effort on trials that demanded greater memory precision. Consistent with our prediction, participants were slower to respond^26–29^ (Figure 1B) and had greater pupil dilation^24,25^ when they anticipated having to make a decision requiring a fine, rather than a coarse, WM representation. Therefore, trying harder had measurable effects on behavior.

We then modeled fMRI activity to characterize how effort impacts the way WM representations are stored in the brain. We first asked how the gain of persistent activity might change with effort. We observed greater amplitude of delay period activity on hard compared to easy trials (Figure 2B). However, the effect of effort on persistent activity was selective rather than equal across the cortex. Effort had the largest effects on the gain of persistent activity in frontal and parietal cortical regions, and little effect in early visual cortex. Increasing memory load, which presumably increases not only difficulty but also effort, also had its greatest effects on BOLD activity in frontal cortex^21,22,30^. Intriguingly, increases in the gain of persistent activity were not restricted to the voxels whose receptive fields contained the memorized target. We used a modified population receptive field (pRF) mapping procedure to measure the receptive fields of voxels in visual field maps not only in visual cortex, but also in frontal and parietal cortex^34^. The gain of persistent activity increased equally across voxels in topographic maps in frontal and parietal cortex. This is important as it suggests that cognitive effort causes a non-selective increase in gain across neural populations. This diffuse increase in gain stands in stark contrast to selective increases in gain associated with spatial attention^38–41^, and therefore suggests distinct neural mechanisms. In the future, a within-subject manipulation of both attentional priority and cognitive effort would allow us to make this comparison directly.

We hypothesized that effort would impact not only the gain of persistent activity, but would improve the quality of WM representations. Recently, we used a Bayesian decoder to estimate both the content and uncertainty of WM representations from delay period activity^35^. Building on these findings, we predicted that increased effort would improve the accuracy and/or decrease the uncertainty of decoded WM representations because both of these decoded metrics predict behavioral WM performance^35,42^. Despite effort impacting the gain of persistent activity in frontal and parietal cortex, we found little evidence that these areas contained information about the stimulus stored in WM. Previous studies have found similar results; areas in association cortex that show robust persistent activity typically lack patterns that can be used to robustly decode the contents of WM^42–44^. Instead, we found that increased effort improved the accuracy of decoded WM representations in early visual cortex and mid-level visual areas that flank the parieto-occipital sulcus (i.e., V3A/B and IPS0) (Figure 3A). On trials when participants needed to exert more effort to maintain a high precision representation of the target, the fidelity of the decoded WM representation in striate and extrastriate visual cortex improved. Decoded accuracy and uncertainty typically co-vary^31,32,35^. However, we found that effort only impacted accuracy, not uncertainty (Figure 3B). It is unclear whether this difference reflects something about how effort impacts WM representations or if it stems from something about our experimental design that did not emphasize memory certainty like our previous study did^35^.

How might our two main findings be related? We hypothesized that the gain of persistent activity in frontal and parietal cortex might be the source of an effort-related feedback signal that targets visual cortical areas where WM targets are stored. Strikingly, we found that trial-by-trial changes in the gain of delay period activity in frontal and parietal cortex predicted trial-by-trial fluctuations in the decoded accuracy of WM targets from the patterns of activity in early visual cortex (Figure 4A). Such a correlation between BOLD activity level and decoding accuracy avoids interpretational ambiguities due to the intrinsic correlations between BOLD activity levels across the brain^45,46^. Moreover, it provides strong evidence in favor of our hypothesis that persistent activity in frontal cortex, which has long been thought to reflect top-down control signals that generally support WM^47,48^, more specifically carry effort-related signals that modulate the quality with which WM representations are stored. The portions of frontal and parietal cortex whose gain predicted WM decoding remarkably overlapped with retinotopically defined maps of space (Figure 4B). Specifically, this included the most caudal part of the PFC in the anterior bank of the precentral sulcus and a swath along the intraparietal sulcus, where previous studies have reported the most robust persistent neural activity during spatial WM^42,49–51^. Notably, perturbations with transcortical magnetic stimulation^52^ (TMS) and surgical resections of the human precentral and intraparietal sulci^53,54^ both cause consistent reductions in the fidelity of spatial WM demonstrating necessity. Perhaps these impacts on WM performance were not due to a direct disruption of the stored WM representation itself, but instead were mediated by impacts on effort-related feedback signals.

Such possibilities highlight the difficulties distinguishing the roles of cognitive effort and the cognitive abilities influenced by such motivational factors, as motivation is rarely systematically controlled. More importantly, this uncertainty has major implications for both understanding the basic mechanisms of the cortical networks that support WM and for understanding the etiology of WM dysfunction in psychiatric disorders. Working memory is both capacity-limited^16,17,55–57^ and costly to recruit^28,58^. Nonetheless, how much variation in WM performance stems from strict capacity limitations versus motivational factors remains unknown. Working memory requires subjective effort to employ^5,58^. People tend to avoid exerting cognitive effort even when they are unaware of doing so or when its avoidance means forfeiting reward^27,28^. That people often avoid recruiting WM suggests that it too may be subjectively costly to use^59–62^. Critically, psychiatric populations appear to suffer from both WM and motivational deficits^63–66^. If at least some of these cognitive deficits, like those in WM, are secondary to the motivational deficits, then attempts should be made to leverage motivational factors to understand psychopathology as well as innovate cognitive remediations.

## Methods

13 participants participated in the study (8 women, mean(sd) age 27(7) years). All participants had normal or corrected-to-normal vision and no history of neurological disorders or brain injury. All participants provided written informed consent in accordance with the Declaration of Helsinki and all study procedures were approved by the Institutional Review Board at New York University. Power analyses based on a comparison to recent fMRI studies with similar aims and methods^21,42^ yielded a goal sample size of 12 participants. One subject was excluded from analyses due to their inability to fixate on the center circle, and the subsequent poor quality of their cortical retinotopic maps (see Methods: Retinotopic Mapping Task). Thus, 12 participants’ data were included in our final sample.

All participants were trained on the task prior to entering the MRI scanner. Inside the scanner, participants made responses with a scanner-safe button box, and their eyes were tracked with Eyelink software. The task was administered using MATLAB Psychtoolbox^67,68^ (The MathWorks, Natick, MA) with screen resolution 1280×1024 and refresh rate 120 Hz. Stimuli were presented using a PROPixx DLP LED projector (VPixx, Saint-Bruno, QC, Canada) located outside the scanner room and projected through a waveguide and onto a translucent screen at the head of the scanner bore. Participants viewed the screen through a mirror attached to the head coil for a maximum viewing distance of 64 cm. Functional image acquisition was locked to the start of the behavioral task through a scanner-sent trigger pulse.

### Behavioral task

Participants completed a single-item spatial WM task (Figure 1A). All participants were instructed on and completed 1-2 practice runs of the task prior to entering the scanner. During the task, participants were required to maintain fixation on a circle at the center of the screen. Fixation was required for the duration of each trial, but not between trials. All stimuli and the fixation circle were presented inside of a dark gray task aperture of 15 degrees of visual angle (dva) in diameter. The fixation circle was light gray, and 0.6 dva in diameter. At the start of each trial the fixation circle turned blue or magenta for 1 second. The color of the fixation circle was indicative of whether the trial at hand would require more or less precise memory of the target’s location. Participants were instructed prior to entering the scanner and again at the start of each run of scanning as to what the color of the fixation circle meant.

The target, a small white dot 0.6 dva in diameter, was presented for 500 milliseconds at a fixed eccentricity of 10 dva. Participants were instructed to maintain the location of the target in memory over a 12 second delay period. During the delay, only the fixation circle and task aperture were visible. At the conclusion of the delay period, a secondary test probe appeared for 800 milliseconds. While the test probe was on screen, participants pressed the “1” key if its location was counterclockwise of the target or the “2” key if it was clockwise. Immediately following this selection, participants were shown the original target and the test probe concurrently. The target reappeared in green if their response was correct, red if incorrect, and yellow if they took too long to respond. Feedback lasted 1 second, and then the inter-trial interval (ITI) was randomly jittered such that it lasted 4.7, 7.7, or 10.7 seconds. During the ITI only the fixation and task aperture remained on screen.

Target locations were pseudorandom such that locations were not repeated, and were always at least 5 degrees away from the horizontal or vertical meridian. Test probes were presented at an angular distance from the target determined by a staircase procedure. This angular distance between target and test was dynamically set for each subject such that accuracy was kept around 70% on trials requiring more memory precision, and 90% on trials requiring less memory precision. Participants accumulated points for correct responses (+1 for every correct choice, +0 for every incorrect choice). At the end of each run, participants were shown the cumulative number of points they had earned that session. Participants completed 12 trials per run and were given a break between runs. Each participant completed up to 13 runs per experimental session for two sessions of approximately an hour and a half each. This yielded a total of 18-20 runs per subject, and 108-120 easy and 108-120 hard trials per subject.

### Retinotopic mapping task

To identify regions of interest (ROIs) for all reported analyses, we acquired retinotopic mapping data based on population receptive field (pRF) modeling^34^. During each retinotopy run, participants completed a rapid serial visual presentation (RSVP) task. Like other retinotopy tasks, ours was designed to collect patterns of BOLD activation while participants experience dynamic stimulation across the entire visual field. Unlike other retinotopy tasks, ours requires that participants not only passively view the screen, but attend and respond to the task. While participants fixated on a center fixation cross, they had to covertly attend to the location of a moving and regularly changing stimulus bar in order to identify rare targets within it. The stimuli were arrays of color photos of objects, which swept across 26.4° of the visual field in twelve 2.6 s steps. Sweep directions were pseudo-randomly chosen from four directions (left-to-right, right-to-left, bottom-to-top, top-to-bottom). Each array was composed of 6 equally sized color images of random household objects. These images were presented for 150 to 600 msec at a time, then replaced with new images. Before each run, participants were shown the target object image for that run, and encouraged to familiarize themselves with it. During the run, participants reported, via a button press, each time this target object appeared in the array of objects. If participants correctly responded within 800 msec of the presentation of the target object, the fixation point briefly turned green. A three-down/one-up staircase was implemented to maintain task difficulty ∼80%, by changing how long one array of 6 objects would be on screen before the next array would replace them. If a subject completed one sweep of the array with less than 80% accuracy, then the object refresh rate was lowered (slowed) for the next sweep. If they were more accurate than 80% in one sweep, then the refresh rate was increased (sped up). The speed of the movement of the array was not changed. Participants were required to fixate on the center of the screen for the duration of the task. Their fixation was confirmed both online and post-data acquisition via gaze tracking.

### Oculomotor methods

We measured gaze and pupil size in the scanner at 500 or 1000 Hz (Eyelink 1000, SR Research, Ontario, Canada), beginning with a five-or nine-point calibration and validation scheme. All 12 participants had at least 1 session of usable eye-tracking and pupil dilation data from within the scanner. Due to technical issues, 1 session of eye tracking was unavailable for 1 participant.

#### Oculomotor pre-processing

To process the eye-tracking data, we used the freely available MATLAB iEye toolbox (github.com/clayspacelab/iEye_ts). We transformed raw gaze positions into degrees of visual angle, removed values outside of the screen area, removed artifacts due to blinks, and smoothed gaze position with a Gaussian kernel centered on 0 degrees of visual angle (dva) with a standard deviation of 5 dva. To improve the accuracy of gaze monitoring, data were corrected for fixation drifts and calibrated to account for measurement noise. Trials were excluded from analyses of pupil size if the participant failed to maintain fixation during the cue, stimulus presentation, or delay periods, or if the quality of the eye-tracking data was poor. These exclusion criteria resulted in removing on average 9% (SD 12%; range 0.4-41.2%) of trials.

#### Pupil size preprocessing

Pupil diameter was extracted from the iEye-processed gaze time courses. To remove scanner-induced high frequency noise from pupil size time courses, we applied a low pass filter with an 8 Hz cutoff. Then, to account for the linear drift in pupil size which generally increases with time spent on task, we subtracted a baseline pupil size from each trial. This baseline pupil size was obtained for each trial by averaging the size of the pupil from seconds 2 to 4 of the preceding ITI. This window was chosen as to not include the pupil dilation associated with the anticipation of the start of the next trial (at the conclusion of the ITI), as well as the tail of the feedback-evoked pupil dilation from the conclusion of the previous trial (at the beginning of the ITI). We also wanted to use a consistent window, even across the variable ITI lengths (of 4.7, 7.7 and 10.7 seconds duration).

To obtain clean pupil time courses and to assess the effect of the difficulty manipulation on pupil dilation, we ran a trial-by-trial regression model on pupil size. The model included an intercept term, the baseline pupil size (during the preceding trial epoch), the X-and Y-coordinates of the subject’s gaze, and the difficulty condition of the trial. We ran three separate regressions within-subject, predicting pupil size 1) only during the stimulus presentation epoch (and the following 1.5 seconds, due to possible sluggishness of the pupil response to light), 2) only during the delay period, and 3) over the time course of the entire trial. Using the beta values obtained from this regression, we constructed a simulated pupil time course, and then subtracted it from the true time course to obtain the residual pupil size time course. We then Z-scored these residuals. We analyzed these Z-scored residuals trial-by-trial, instead of the raw pupil size data, as a way of removing the confounding influence of gaze position and baseline pupil dilation on pupil size. In order to visualize the mean effect of the difficulty manipulation on pupil dilation, we did not use the difficulty predictor value in the pupil time course reconstruction, using only the intercept, baseline, and eye position terms.

### fMRI methods

#### Anatomical and bias images

T1-and T2-weighted images were acquired using the Siemens MPRAGE and Turbo Spin-Echo sequences (both 3D) with 0.8 mm isotropic voxels, 256 × 240 mm slice FOV, and TE/TR of 2.24/2400 ms (T1w) and 564/3200 ms (T2w). We collected 192 and 224 slices for the T1w and T2w images, respectively. We acquired between two and four T1w images per subject and one to three T2w images, which were aligned and averaged to improve signal-to-noise.

#### fMRI acquisition

All functional MRI images were acquired at the NYU Center for Brain Imaging on a 3T Siemens Prisma Scanner. The experimental and retinotopic mapping task were imaged using the CMRR MultiBand Accelerated EPI Pulse Sequences^69–71^. All functional and anatomical images were acquired with the Siemens 64 channel head/neck coil. All scanning sessions started with two fast 3D GRE sagittal images (resolution: 2 mm isotropic, FoV: 256 × 256 × 176 mm; TE/TR: 1.03/250 ms), one with the body coil and the other with the 64 ch head/neck coil. We collected these images in order to correct functional images for inhomogeneities in the receive coil sensitivity and improve the motion correction and coregistration process during pre-processing.

For the experimental scans, BOLD contrast images were acquired using a Multiband (MB) 2D GE-EPI with MB factor of 4, 44 2.5-mm interleaved slices with no gap, isotropic voxel size 2.5 mm and TE/TR: 30/750 ms. We measured field inhomogeneities by acquiring spin-echo images with normal and reversed phase encoding (3 volumes each), using a 2D SE-EPI with readout matching that of the GE-EPI and same number of slices, no slice acceleration, TE/TR: 45.6/3537 ms.

For the retinotopic mapping scans, BOLD contrast images were acquired using a Multiband (MB) 2D GE-EPI with MB factor of 4, 56 2 mm interleaved slices with no gap, isotropic voxel size 2 mm and TE/TR: 42/1300 ms. Distortion mapping scans were acquired with normal and reversed phase encoding, using a 2D SE-EPI with readout matching that of the GE-EPI and same number of slices, no slice acceleration, TE/TR: 71.8/6690 ms.

During both experimental and retinotopic mapping scans, we divided functional sessions into 2–5 ‘mini-sessions’ consisting of 1–5 task runs split by a pair of spin-echo images acquired in opposite phase encoding directions. We repeated this spin-echo imaging within-scanning session because the distortion field can depend on the exact position of the head within the main field, which sometimes slowly changes over the course of the session. We then used these spin-echo images for anatomical registration and for distortion correction for all functional MRI scans collected.

#### fMRI preprocessing

We used all intensity-normalized high-resolution anatomical scans as input to the ‘hi-res’ mode of Freesurfer’s recon-all script (version 6.0) to identify pial and white matter surfaces. We edited these surfaces by hand using Freeview as necessary and converted surfaces to SUMA format. The processed anatomical image for each participant acted as the alignment target for all functional data. Our aim for preprocessing was to put functional data from each run into the same space at the same voxel size acquired during the task sessions (2.5 mm isovoxel), account for run-and session-specific distortions, incur minimal volume-wise smoothing by minimizing spatial transformations, and apply a marginal amount of smoothing along the direction orthogonal to the cortical surface. This allowed us to optimize SNR and ensure that functional data remained as near as possible to its original 2D location on the cortical surface.

Recall that we recollected spin-echo images every 1–5 task runs, resulting in 2-5 mini task sessions. We applied all preprocessing steps described below to each mini-session independently. Preprocessing was performed using a combination of scripts generated with AFNI afni_proc.py and custom scripts implementing AFNI functions (version 17.3.09, pre-compiled Ubuntu 16 64-bit distribution). We performed all analyses on a LINUX workstation running Ubuntu v16.04.1 using 8 cores for most OpenMP accelerated functions.

During preprocessing, we first corrected functional images for intensity inhomogeneity induced by the high-density receive coil by dividing all images by a smoothed bias field (15 mm FWHM), computed as the ratio of signal in the receive field image acquired using the head coil to that acquired using the in-bore body coil. To improve coregistration of functional data to the target T1 anatomical image, we used distortion-corrected and averaged spin-echo images (which were used to compute distortion fields restricted to the phase encoding direction) to compute transformation matrices between functional and anatomical images. Then, we used the rigorous distortion correction procedure implemented in afni_proc.py to undistort and motion correct functional images. Briefly, this procedure involved first distortion correcting all images in each run using the distortion field computed from the spin-echo image pair, then computing motion correction parameters (6-parameter affine transform) using these unwarped images. Next, we used the distortion field, motion correction transforms for each volume, and the functional-to-anatomical coregistration simultaneously to render functional data from native acquisition space into unwarped, motion corrected, and coregistered anatomical space for each participant at the same voxel size as data acquisition in a single transformation and resampling step. For retinotopic mapping data, this was a 2 mm isovoxel grid, and for task data, this was 2.5 mm isovoxel grid. For both task and retinotopy data, we projected this volume-space data onto the reconstructed cortical surface. For retinotopy data, we made a smoothed version of the data by smoothing on the cortical surface (5 mm FWHM). We then projected surface data (for task data, only the ‘raw’ data; for retinotopy data, the raw and smoothed data) back into volume space for analysis. For unsmoothed (task) data, this results in a small amount of smoothing for each voxel along a vector orthogonal to the surface.

To compute pRF properties (see below) in the same voxel grid as task data, we projected retinotopy time series data onto the surface from its native space (2 mm iso), then from the surface to volume space at the task voxel resolution (2.5 mm iso). We used this volumetric 2.5 mm data to fit the pRF model. This ensured that variance explained estimates faithfully reflect goodness of fit and are not impacted by smoothing incurred from transforming fit parameter values between different voxel grids. It also ensures accurate translation of ROIs drawn based on pRF model fits to functional data obtained during the experimental task.

### Data analysis

#### Behavioral and oculomotor data analysis

To confirm the success of our staircase procedure, we computed the mean response accuracy across “easy” and “hard” trials for all participants. We also computed the mean response times across task conditions. We ran planned t-tests to compare the mean accuracy and response times (RTs) across trial types.

To assess the time course of effort-related pupil dilation during the WM delay period, we ran a sliding window procedure comparing mean pupil size across hard and easy trials on the group level. Each window was 50 milliseconds in length (25 samples per condition). We ran a t-test over each window comparing the distributions of mean pupil dilations for hard and easy trials, with data aggregated across all 12 participants. As we tested 306 separate time windows (assessing both the stimulus presentation and delay periods), we corrected for multiple comparisons by assessing significance at a Bonferroni-corrected level of 0.00016. We then plotted the significance of timepoints during the delay period by overlaying stars at the center of each significant window (Figure 1D).

Within-subject, we assessed the significance of the effect of the difficulty manipulation on pupil dilation by assessing the confidence intervals around the predictor betas, and whether they included 0. Most participants’ pupils were significantly more dilated during the delay period of hard trials compared to easy trials (within-subject trial-by-trial regression analyses yielded significant difficulty terms in 9/12 participants). The difficulty manipulation significantly increased pupil dilation in the stimulus presentation period for 7/12 participants, and over the course of the entire trial in 7/12 participants. All results from the fMRI data were the same when we included only the 9 participants with significant pupil differences.

#### pRF Model Fitting and ROI drawing

We averaged time series from each voxel across all retinotopy runs (9-12 per participant) in volume space and fit a pRF model for each voxel using a GPU-accelerated extension of vistasoft (github.com/clayspacelab/vistasoft). We fit a compressive spatial summation isotropic Gaussian model as implemented in mrVista (see ^34^ for a detailed description of the model). We created a high-resolution stimulus mask (270 × 270 pixels) to ensure similar predicted response within each bar size across all visual field positions (to mitigate the effects of aliasing with a lower-resolution stimulus mask grid), and began with an initial high-density grid search, followed by subsequent nonlinear optimization. Note that, in these analyses, because we conducted a grid search on all voxels independently, there was no spatial smoothing of parameter estimates across neighboring voxels prior to the nonlinear parameter search. For all analyses described below, we used best-fit pRF parameters from this nonlinear optimization step.

After estimating pRF parameters for every voxel in the brain, ROIs were delineated by projecting pRF best-fit polar angle and eccentricity parameters with variance explained ≥10% onto each participant’s inflated surfaces via AFNI and SUMA. ROIs were drawn on the surface based on established criteria for polar angle reversals and foveal representations^34,72–74^. Finally, ROIs were projected back into volume space to select voxels for analysis. In this report we considered data from V1, V2, V3, V3AB, intraparietal sulcus (IPS) area IPS0, IPS1, IPS2, IPS3, inferior precentral sulcus (iPCS), and superior PCS (sPCS). For simplicity, we combined data across ROIs based on shared foveal representations, resulting in larger ROIs for analysis: V1-V3, V3AB, IPS0-IPS1, IPS2-IPS3, iPCS, and sPCS. Only voxels with ≥10% variance explained in pRF model fits were included in subsequent fMRI analyses.

In addition to using these pRF parameters to draw functional ROIs in surface space, we used them to select specific voxels for analysis trial-by-trial. The pRF model provided an estimate of the location and size of each voxel’s receptive field (RF). Using the PRF-fit parameters for each voxel, we drew simple binary receptive fields containing the fit center of the RF and 15 degrees on either side. If the stimulus location fell within that voxel-specific box, then that voxel’s activity was labeled “in RF” on that trial. Voxels with receptive fields that contained the point 180 degrees away from the stimulus (or points ≤15 degrees away) were labeled “out RF”. We performed univariate analyses of delay period BOLD responses comparing across in RF and out RF voxels (“RF type” analyses).

#### Univariate fMRI analyses

To visualize the effect of task difficulty on average BOLD activity, we computed percent signal change of the BOLD signal, using the BOLD activity on the first 3 TRs of each trial as the denominator. Using the beginning of the trial as the baseline also linearly detrended the BOLD data. We computed percent signal change separately for each voxel within each ROI, then averaged across voxels, trials, and participants to produce average time courses within our conditions of interest. We then plotted the group mean time courses for each ROI. Standard error of the mean (SEM) was calculated across participants’ average time courses by taking the standard deviation across participants, then dividing by the square root of the sample size (N=12). SEM was then used to draw error bars. To assess differences between conditions, voxel types and ROIs, we performed a 3-way repeated measures ANOVA on late delay period activity (seconds 6-12 of the delay). We confirmed the significance of the F-values obtained by this ANOVA with permutation testing. We constructed null distributions of F values by shuffling the condition, ROI, and RF type labels 1000 different times, assessing the F-statistic from that shuffled dataset each time. We then compared the original F-statistic to the null distribution of F-statistics obtained by shuffling, and obtained a p-value from the number of F-statistics which were greater than or equal to the original F. Interactions between variables were investigated using post-hoc t-tests within ROIs, comparing across condition means (RF type and trial difficulty).

We trained and subsequently tested the Bayesian decoder (TAKFAP; see: Bayesian Decoding with TAKFAP) using the mean percent signal change of each voxel in each ROI from seconds 6 to 12 of the delay period, yielding one number per voxel per trial.

#### Trial-by-trial general linear model

We estimated the magnitude of BOLD responses during the WM delay period (the delay period magnitude) with a general linear model (GLM) which included separate regressors for each trial. We created the trial-wise model in AFNI using *3dDeconvolve*. For 10 participants, this resulted in 240 delay period regressors. Two participants completed only 228 and 216 trials, resulting in fewer regressors. Each regressor modeled the activity evoked during the delay of one single trial using a boxcar function, time-locked to delay period onset and convolved with an assumed hemodynamic response function (Gamma function) using the ‘GAM(p,q,d)’ function in AFNI. The parameters of the Gamma function were set to the default values (p=8.6, q=0.547), except for the duration (d) parameter, which was set to match the length of the delay period (12 seconds).

The GLM also included two regressors to account for average cue and response period activity and six motion regressors obtained using afni_proc.py during the motion correction step of preprocessing. The ordinary least squares solution to this trial-wise GLM yielded a unique activation estimate (β value) for each delay period.

#### Bayesian decoding with TAFKAP

We obtained trial-by-trial estimates of WM representational accuracy and uncertainty using a Bayesian decoder^31,32^. To implement this decoder, we used the publicly available TAFKAP toolbox (https://github.com/jeheelab/TAFKAP). The TAFKAP method provided trial-by-trial probability distributions over possible memorized stimulus values based on the patterns of BOLD activation on those trials. Specifically, in training the algorithm learned a generative model relating trial-by-trial voxel-wise activation patterns to the true location of the target stimulus on those trials. Then it inverted the learned associations between patterns of activation and stimulus locations to provide the posterior probability of the target stimulus’s location, centered on the mean (maximum likelihood) value:

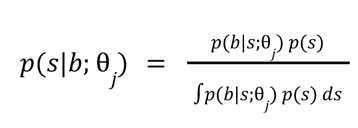

Where *b* is the multivariate voxel response to each stimulus *s*, and θ is the set of free parameters which were trained on one set of training data *j.* To determine the likelihood of voxel activation patterns *b* given the stimulus on screen *s*, the decoder assumed that each voxel had a tuning curve which could be approximated as a weighted sum of eight basis functions which evenly tile location space. The basis functions were half-wave rectified sinusoidal functions raised to the power of 8 and centered on evenly spaced means φ :

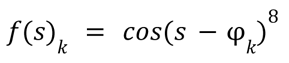

The response of each voxel *b_j_* given stimulus *s* was then modeled as a weighted combination of these basis functions’ responses to *s*:

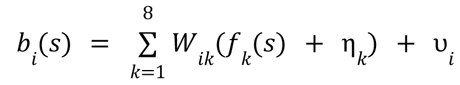

The weighting matrix W determined the degree that the tuning and noise function from each basis function contributed to the response of each voxel. The noise term of each basis function η was applied to the same extent as the activation function of that basis *f*(*S_k_*), capturing that similarly-tuned voxels exhibit similar noise. *υ_j_* was the noise specific to voxel *i*. TAFKAP assumes that both sources of noise are normally distributed with mean 0. Fitting these individual noise terms for each voxel may not be completely possible, especially if the number of voxels to fit is larger than the number of trials in the dataset. Thus TAFKAP shrinks the sample covariance matrix to a simpler target covariance matrix, to a degree determined by free parameter λ. More details of this fitting procedure are available in ^32^.

Our input to the model was the z-scored mean percent signal change during the late delay period (6-12 seconds of delay) for voxels within each ROI. Late delay period activity was z-scored within-run. We ran the model separately for each of our ROIs. We specified a flat prior *p(s)* over all possible target stimulus locations, as the polar angles of the target stimuli were drawn from a uniform distribution. Each set of training data was bootstrap resampled to improve the fidelity of the recovery process. For each trial in the test data, the decoding results from each resampling were averaged to obtain one decoded posterior probability distribution:

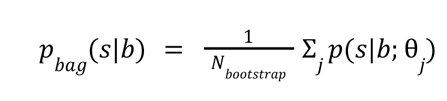

Then, a trial-specific estimate of the remembered stimulus location was obtained by taking the circular mean of the posterior inferred from the mean voxel-wise activity on that trial. The uncertainty around the estimated location was obtained by taking the circular standard deviation of the posterior. The TAFKAP algorithm has been used to relate representational accuracy and uncertainty to behavioral variability and confidence reports during perception and WM^31,35^. Thus there is evidence that TAFKAP reliably measures the quality of WM representations.

To test whether representational quality changed according to task demands, we performed a 2-way repeated measures ANOVA on WM decoding accuracy and uncertainty with task condition and ROI as the main factors. If we obtained a main effect of trial type or ROI, we ran permutation testing. The true F-scores for each main effect were compared to a null distribution of F-scores obtained by shuffling trial type and ROI labels 1000 times each. P-values were determined by computing the number of F-scores greater than or equal to the true F-score. In addition, we ran t-tests to assess condition differences in representational quality within each ROI.

In addition to understanding differences in condition effects in our ROIs, we aimed to quantify which ROIs contain decodable stimulus information during the delay period. To assess this, we ran circular correlations of decoded stimulus positions versus true stimulus positions trial-by-trial. We then compared the group mean circular correlation values against 0 using a t-test (Supplementary Figure 1).

#### Correlation between delay period magnitude and TAFKAP outputs

Here and in other work^42,43^ we observed a tradeoff across ROIs between decoding accuracy and the magnitude of delay period activity. For example, sPCS demonstrated low decoding accuracy but high activation above baseline during the WM delay. To shed light on the distinct roles of and possible interactions between neural regions, we performed a trial-wise correlation analysis between decoding in visual cortex and amplitude of delay period activity in all voxels in the brain. Our goal was to test the hypothesis that the quality of WM representations in visual cortex were predicted by the magnitude of delay period activity in association cortex. To do so, we correlated delay period betas (magnitudes) with decoded error and uncertainty from visual cortex, specifically V1-V2-V3. With our Bayesian decoder, we obtained trial-wise probability distributions of the location of the memoranda, as a measure of the quality of WM representations. The memory error and uncertainty were derived from the mean and standard deviation of the posterior, respectively (see *Bayesian decoding with TAFKAP)*. Note that the predicted relationship between delay period activity and decoding was likely present during both easy and hard trials. Thus, to maximize our power to detect the predicted correlation, we computed correlations across all trials. Future studies with more power from more trials might be able to test how task difficulty might mediate these correlations.

In order to interpret the correlation magnitudes, we obtained within-subject z-scores by applying an inverse hyperbolic tangent transform to the correlation coefficients, then used these for group-level testing and visualization. To reduce noise on the z-score maps, we applied a 5-mm full-width at half-maximum (FWHM) Gaussian kernel for spatial smoothing using AFNI’s *3dmerge* function. To make map-based inferences at the group level, the z-transformed correlation maps of individual participants were spatially normalized from their native space into the normalized Montreal Neurological Institute (MNI 152) space. We first obtained each subject’s high-resolution anatomical image (0.8 mm isotropic voxel), aligning its orientation and voxel size with the standard MNI 152 template (MNI152_T1_2009c) using *3dresample*. We then registered the anatomical images to the MNI template using the automatic Talairach (affine) transformation function (*@auto_tlrc*). With the output registration parameters obtained from registration, we then use *3dAllineate* to transform individual z-maps into standard MNI coordinates.

To assess the significance of our effects on a group level, we performed a t-test using AFNI’s 3dttest++ function to identify voxels for which the mean of transformed and smoothed correlation coefficients across participants was significantly greater than zero. Once we obtained an estimate of which voxels showed significant effects, we mitigated the increased false alarm rate associated with running multiple comparisons by running a cluster permutation test to identify significant clusters, not voxels. We conducted a cluster-based simulation to estimate the probability of false positive clusters using *3dClustSim* in AFNI. First, we calculated the smoothness of noise from our subject-level maps, using the *3dFWHMx* function on each individual’s residual time series (errts files from AFNI *3dDeconvolve*), and obtained subject level smoothness in three dimensions (x, y, z). Next, we determined the cluster size threshold using *3dClustSim* (input averaged smoothness across participants, see Supplementary Table 1 for simulation results) and displayed the group-level results using the correct thresholding (Figure 5A).

After obtaining significant clusters at the group level, we wanted to verify that they were overlapping with our functionally-defined regions of interest. Because each subject’s ROIs were drawn in native, not template space, we first extracted each subject’s ROIs (see *Retinotopic mapping and ROI definition)* and transformed them into MNI template space. We then created an overlay in template space of all individuals’ ROI masks, to create a group-level functional ROI map. Each voxel in the overlay had a value from 0 to 12, indicating how many participants’ ROIs contained that voxel (Figure 5B). This map allowed for visual inspection of our significant clusters and the location of the original functional ROIs, to confirm their co-location.

## Data & Code Availability

All analysis code used in this work is available on a public GitHub repository: https://github.com/clayspace/MasterLiCurtis2024. The raw, unprocessed fMRI data and the functionally-defined regions of interest analyzed here are available in NIFTI format on the Open Science Framework: https://osf.io/c6yjg/.

## Acknowledgments

We gratefully acknowledge Hsin-Hung Li, Grace Hallenbeck, and Yuna Kwak for their support during fMRI data collection, pre-processing, and analysis. We thank Kartik Sreenivasan for his thoughtful input on our neuroimaging analyses. We thank Keith Sanzenbach, Chrysa Papadaniil, and the staff at the Center for Brain Imaging at NYU for their assistance designing and testing fMRI protocols. Lastly, we thank the rest of the Curtis Lab for many helpful conversations about this work. This research was supported by the National Eye Institute (R01 EY-016407, R01 EY027925 and R01 EY033925 to CEC), the NYU MacCracken Fellowship (to SLM), and the NIH (Vision Training Grant 5T32EY007136-29 to SLM).

## Supplementary Information

**Supplementary Figure 1:**
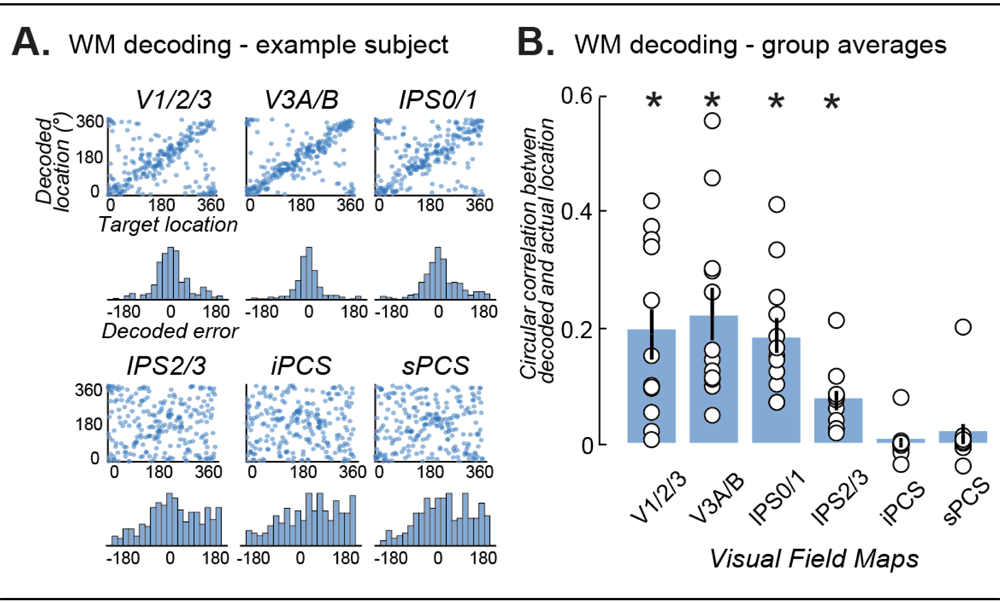
Decoding fidelity on the individual and group levels. **A.** Decoding fidelity in each ROI for one subject. A Bayesian decoder (TAFKAP) was applied to mean late delay period (seconds 6-12) activations as predictors and true stimulus locations as labels, separately for each ROI. Stimulus polar angle in degrees was decoded with a leave-one-run-out cross-validation procedure trained on 16-18 runs (196-216 trials) and tested on 2 runs (24 trials) of data from each participant. We implemented 9 to 10 training/test folds. This entire procedure was then iterated over. True target positions (in polar angle) are plotted on the X-axis, and decoded target positions are plotted on the Y-axis. Lower visual areas V1-V3 and V3AB and parietal area IPS0-IPS1 exhibit very high decoding accuracy, while parietal area IPS2-IPS3 and frontal areas iPCS and sPCS exhibit lower decoding accuracy. **B.** Mean group-level fidelity of the Bayesian decoder in recovering the true stimulus locations from mean BOLD activity during the late delay period (seconds 6 to 12). Circular correlation between the true stimulus location and decoded stimulus location was obtained for each subject for each ROI. Individual values are plotted in filled circles. Group means are plotted in black. Errorbars were obtained using the standard error of the mean. Each ROI’s decoding fidelity was assessed with a t-test comparing the group circular correlation values to 0. Stars indicate significance at a level of p < 0.05.

**Supplementary Figure 2:**
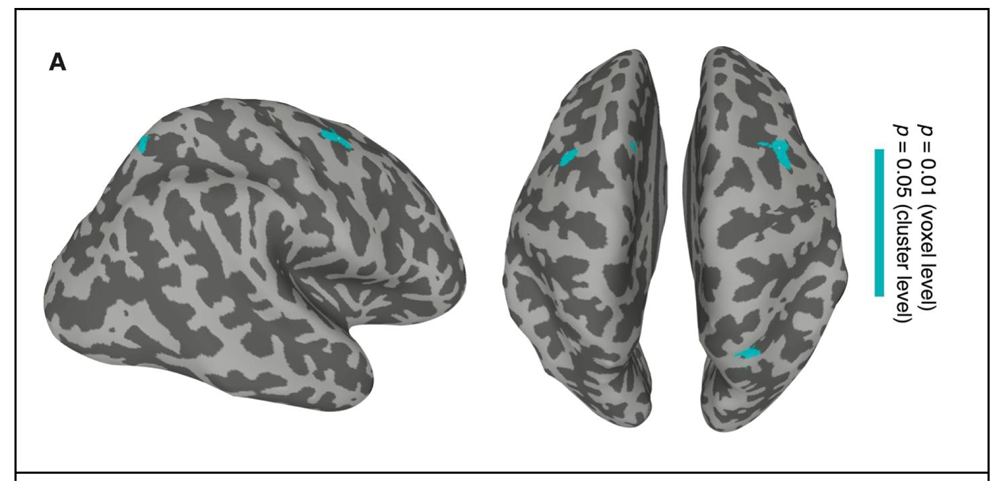
Gain of delay period activity predicts decoded uncertainty from early visual cortex. **A.** To identify potential sources of top-down control over representational fidelity in visual cortex, we created a whole-brain correlation map. Trial-by-trial magnitudes of delay period activity across the whole brain were the predictors. The predicted term was trial-by-trial decoded uncertainty from V1-V2-V3. The cyan clusters are areas with significant negative correlation at the group-level using cluster-based permutation testing. A negative correlation with decoded uncertainty indicates a positive relationship between representational quality in visual cortex and delay period magnitude in the cortical regions which are filled in in cyan.

**Supplementary Table 1:** Permutation result table. Each row indicates the corresponding voxel level p value, each column indicates corresponding cluster level alpha value.

| $\alpha$ | 0.1 | 0.05 | 0.02 | 0.01 |
| --- | --- | --- | --- | --- |
| $p$ threshold | | | | |
| 0.05 | 1680.5 | 1904.0 | 2242.0 | 2416.0 |
| 0.02 | 539.5 | 611.6 | 698.5 | 766.0 |
| 0.01 | 299.7 | 340.6 | 390.5 | 427.0 |
| 0.005 | 182.8 | 208.7 | 245.0 | 267.5 |
| 0.002 | 103.9 | 119.8 | 140.8 | 156.6 |
| 0.001 | 71.1 | 82.0 | 97.6 | 109.7 |
| 0.0005 | 49.9 | 57.8 | 69.9 | 80.0 |
| 0.0002 | 31.9 | 37.8 | 46.7 | 52.8 |
| 0.0001 | 23.2 | 27.8 | 35.2 | 39.9 |

